# Geometry-based dynamics of the postsynaptic density explain protein capture by an actin-spine-geometry-dependent synaptic tag

**DOI:** 10.64898/2026.07.16.738887

**Authors:** Mitha Thomas, Michael Fauth

## Abstract

The synaptic tagging and capture (STC) hypothesis explains how early-phase plasticity is converted into its late phase through the coincidence of synaptic tagging and plasticity-related protein (PRP) availability. Yet the biophysical basis of this process remains poorly understood. Based on the hypothesis that the interaction of actin and spine geometry implement the synaptic tag, we here investigate the associated PRP capture mechanism. We propose that capture is implemented by PSD remodelling which is gated by local membrane curvature at the PSD periphery. Using computational modelling, we show that curvature variations around the PSD that arise from long-term potentiation (LTP) inducing stimuli indeed enable a PSD growth, reproducing late-phase potentiation and the maintenance of structural LTP. We further explore how the timing of PRP availability relative to tag formation and the initial spine size determine the extent of PSD enlargement, yielding outcomes consistent with experimental findings. Hence, our results support a structural interpretation of synaptic tagging and capture in which a transient, actin-driven geometric state of the spine encodes the tag, and curvature-mediated PRP recruitment stabilises synaptic changes, and thus render spine geometry as a key biophysical regulator of memory consolidation.

## 1 Introduction

The cellular basis of memory formation in the brain is believed to be implemented by changes in the strength of the synapses that connect neurons with each other, broadly termed as synaptic plasticity. Ever since its discovery in 1973, long term potentiation (LTP) (Bliss and Lømo, 1973), characterised by an increase of synaptic transmission efficacy, has been a standard model of synaptic plasticity. LTP occurs in two phases: the early phase (e-LTP) that constitutes the initial expression of LTP, and the late phase (l-LTP) which sustains it at least over several hours (Huang, 1998). The synaptic tagging and capture hypothesis (Frey and Morris, 1997) explains that for the emergence of l-LTP, the stimulation sets a transient synaptic tag at activated synapses, enabling them to subsequently capture newly synthesised plasticity-related products (PRPs) at their postsynaptic densities (PSDs). Through this process, initially labile synaptic changes can be stabilised into long-lasting modifications, a phenomenon commonly referred to as synaptic consolidation (Clopath, 2012). Classical STC experiments (Frey and Morris, 1997, 1998; Martin and Kosik, 2002; Redondo and Morris, 2011) have demonstrated that tag setting and PRP synthesis can be independent processes. In particular, the weak-before-strong (WBS) stimulation protocol (Frey and Morris, 1998) shows that synapse stimulated by a weak tetanus do not elicit PRP synthesis but still become tagged which allows them to capture PRPs synthesised as a result of a strong stimulus at another synapse as long as the tag persists. These experiments have identified several defining properties of the synaptic tag (Martin and Kosik, 2002; Redondo and Morris, 2011): (i) synapse specificity, whereby only stimulated synapses become tagged; (ii) tag life- time, which persists on the order of 1–3 hours; (iii) independence from protein synthesis, whereby the tag formation by a stimulus does not require PRP synthesis; and (iv) the ability to capture PRPs to stabilise potentiation.

Several candidates have been proposed to act as the tag, mostly from a biochemical perspective highlighting different molecules, a few among which are CaMKII (Martin and Kosik, 2002; Okamoto et al., 2009, 2007; Redondo et al., 2010), PKA (Huang et al., 2006; Park, 2024), PKM*ζ* (Smolen et al., 2012; Sajikumar et al., 2005), and the cytoskeletal protein actin (Pinho et al., 2020; Szabó et al., 2016). Yet, no single molecule could fully explain the synaptic tag and its precise identity remains an elusive question (Redondo and Morris, 2011). Recent revisions of the STC hypothesis describe the synaptic tag as a structural state of the synapse potentially involving many protein interactions as well as morphological remodelling of the spine and its internal organisation, essentially acting as a mechanical substrate for facilitating long term synaptic changes (Redondo and Morris, 2011).

The remodelling of dendritic spines is strongly linked to the scaffolding protein actin, a candidate already highlighted in STC. Actin forms filaments that organize into networks which contributes to spontaneous spine shape fluctuations under basal conditions (Fischer et al., 1998; Bonilla-Quintana et al., 2020) as well as activity-dependent morphological changes following synaptic stimulation (Bosch et al., 2014; Thomas et al., 2025). The biophysics of actin-driven morphological changes of cellular membranes is explained by the Brownian ratchet theory (Peskin et al., 1993; Mogilner and Oster, 1996), in which thermal fluctuations of the membrane are rectified by actin polymerisation to produce directed membrane deformation. The polymerisation dynamics of actin filaments and their organization into networks is regulated by a variety of actin binding proteins (ABPs, Blanchoin et al., 2000; Borovac et al., 2018). Actin in spines also exhibits heterogeneous dynamics (Frost et al., 2010) and exists in at least two pools: a dynamic pool, constituted by rapidly polymerising filament networks with a turnover rate in the order of a few seconds to one minute, and the stable pool, constituted by filaments with a relatively slower turnover rate, in the order of 17 minutes (Star et al., 2002; Frost et al., 2010; Honkura et al., 2008). Upon LTP, both these pools undergo significant remodelling through variations in the concentrations of ABPs (Bosch et al., 2014). This is accompanied by an enlargement of the spine (Matsuzaki et al., 2004; Okamoto et al., 2009), commonly known as structural LTP (sLTP). The tight link between these processes has led to the proposal of actin and spine geometry as a potential tagging mechanism (Redondo and Morris, 2011; Pinho et al., 2020; Okuda et al., 2021; Thomas et al., 2025; Negri et al., 2025). Using a combined approach of modelling and FRAP experiments, we recently showed that LTP-induced spine geometry and actin alterations indeed persist on the timescale property of the synaptic tag (Thomas et al., 2025). Moreover, the other defining properties of the synaptic tag described above, namely synapse specificity and independence from protein synthesis can be assumed to be fulfilled for spine geometry and filamentous actin. The focus of the present study is therefore to investigate the remaining property and clarify *whether and how actin and spine geometry based synaptic tag can lead to the capture of PRPs*.

We argue that PRP capture is implemented by the growth of the postsynaptic density (PSD), which can be gated by the membrane geometry around its periphery, and therefore depends on the above proposed synaptic tag. To provide insights on how such a tagging and capture mechanism influences long-term potentiation, we investigate a computational model with a PSD-dynamic based on the membrane geometry, which assumes that the smaller or even inverted membrane curvature around the PSD enables PSD growth and, thus, PRP capture. We demonstrate that this geometry-based mechanism is sufficient to enable sustained PSD growth after LTP, and further characterise its operating regime. By testing how the timing of PRP availability relative to tag formation influences the extent of PSD enlargement, we reproduce the distinct plasticity outcomes observed in the weak-before-strong stimulation protocol(Frey and Morris, 1998). In addition, we find that the initial size of dendritic spines modulates plasticity outcomes, which matches experimental findings (Meyer et al., 2014; Bosch et al., 2014; Sun et al., 2021). Taken together, these results demonstrate the viability of a geometry-based synaptic tagging and PRP capture mechanism.

## 2 Results

### Membrane geometry controls PRP capture at the PSD

The capture of LTP-related PRPs elicits long-lasting LTP – that is an increase of the excitatory postsynaptic potentials (EPSPs) at the respective synapse. Under basal conditions these EPSPs correlate with the PSD size through the number of anchored AMPA receptors (Holler et al., 2021; Marrone and Petit, 2002; Becker and Tetzlaff, 2021). Hence, to elicit long-lasting EPSP growth, PRP capture should cause the growth of the PSD. PSD growth after LTP is usually delayed by 15-30 minutes or more (Meyer et al., 2014; Bosch et al., 2014; Hayashi et al., 2009) which also matches with the necessity to synthesize new proteins. Moreover, protein synthesis blockers can negate the growth of the PSD as measured by the incorporation of PSD-scaffolding-proteins such as PSD95, Homer and Shank (Bosch et al., 2014), indicating that PRP availability is a necessary condition for LTP-inudced PSD growth. This is further supported by the fact that Homer1 itself is considered a PRP candidate (Prodan and Morris, 2024) and other PRP candidates like PKM*ζ* bind and cluster the above mentioned PSD molecules (Shao et al., 2012). Thus, in this study we assume that the relevant mechanics of PRP capture in LTP occurs through the growth of the postsynaptic density by integrating the newly synthesized PRPs (compare Redondo and Morris, 2011).

The PSD is typically structured into multiple layers (Fig. 1A, Dosemeci et al., 2016; Zeng et al., 2018; Chen et al., 2008). First, a membrane layer which comprises membrane bound proteins such as glutamate receptors or cell adhesion molecules like neuroligin. This layer is followed by a core layer in which PSD95 molecules extend orthogonal to the membrane (Chen et al., 2011) and then bind with scaffolding proteins Homer and Shank in the pallium layer. These proteins form a preserved, mesh-like polymeric structure with mesh-sizes around 20 *−* 40 nm (Blanpied et al., 2008; Hayashi et al., 2009). In order to preserve this layered structure, PSD can be assumed to grow only parallel to the layers and thus at its outer boundary (Fig 1B). The spine membrane at this boundary can then influence the PSD growth. In particular, if the membrane tightly wraps around the PSD, there is no space to extend the scaffolding mesh (Fig. 1E, condition 3), but if the membrane runs parallel to the PSD layers or even bends away, the PSD can grow (Fig. 1E, conditions 1&2). Biophysically, this can be described by a Brownian ratchet mechanism (Peskin et al., 1993; Mogilner and Oster, 1996). In this framework, the thermal membrane fluctuations must be large enough to eventually leave enough space to integrate new proteins into the mesh. For a linear polymer filament, the probability to add a monomer is attenuated by exp 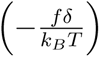 (Peskin et al., 1993), where *δ* is the increase in filament length through adding a monomer, *f* is the membrane counterforce and *k_B_T* the thermal energy. For actin, with *δ ∼* 2.5nm, typical membrane forces only slow down the polymerization. For extending the PSD mesh, *δ* is tenfold larger, which translates into the growth probability exponentially such that typical membrane forces likely stall PSD growth. However, following LTP, spines expand strongly which reduces the membrane counterforce or even bends the membrane away from the PSD boundary, such that growth is enabled. In this way, spine geometry alterations could gate the growth of the PSD due to PRP capture.

**Figure 1:**
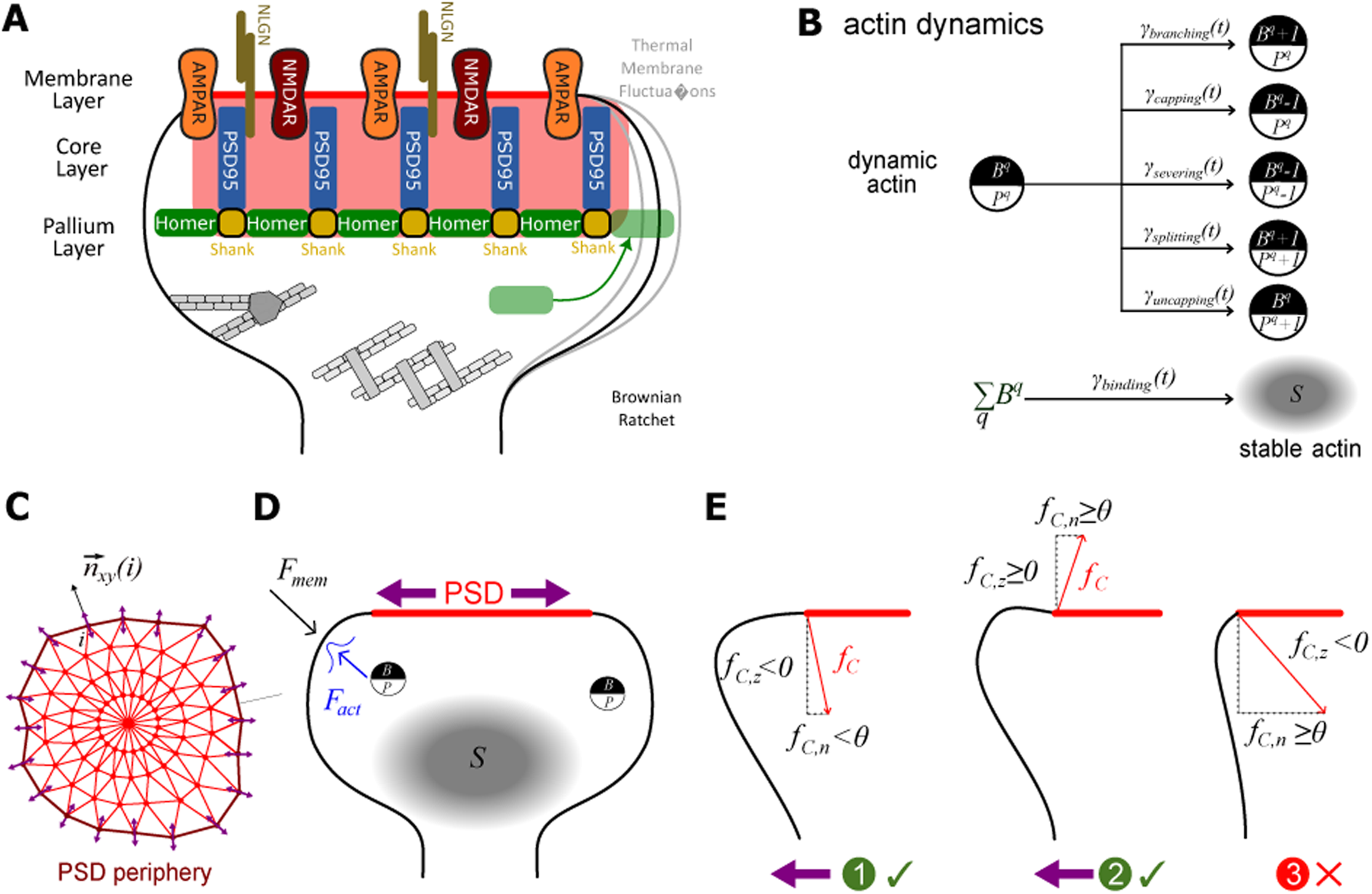
Model overview (A) Schematic of the layered structure of the PSD (red shaded) with the proposed ratchet mechanism for PSD growth: thermal fluctuations must make enough space to insert new PSD scaffolding proteins (B) Actin dynamics at a certain focus *q* based on different ABP-associated processes. Amount of dynamic actin (number of barbed ends *B* and pointed ends *P*) stochastically evolves based on the rates (*γ*(*t*)) of each process. Amount of stable actin (*S*) evolves from the total amount of dynamic actin through crosslinker-associated binding and unbinding rates. (C) Top view of the PSD mesh points. At each point, PSD dynamic is implemented by movement along the normals within the PSD-plane (x-y-plane). (D) Spine with dynamic and stable actin, and PSD represented as a disc (red). *F_act_* is the force generated by actin polymerisation and pushes the membrane outward, while *F_mem_* is the membrane counterforce. (E) PSD dynamics governed by the curvature-induced component (*f_C_*) of *F_mem_*. Conditions 1 and 2 permit PSD growth, whereas condition 3 does not.

### Mathematical Model

To test whether this intuition of geometry-dependent gating of protein capture is viable, we here develop a mathematical model to simulate spine geometry and PSD dynamics during LTP. To this end, we extend our existing spine-geometry model (Bonilla-Quintana et al., 2020; Thomas et al., 2025), which is shortly summarized in the following: The dendritic spine is represented by a discretised mesh, with a fixed subset of points for the spine neck and a subset of points in one plane representing the PSD (red lines in Fig. 1D-E, top projection in Fig. 1C). We assume the dynamic actin pool to consist of multiple actin polymerisation foci (Frost et al. (2010), black & white circles in Fig. 1B&D). Each focus *q* is described by their number of barbed filament ends *B^q^* and pointed filament ends *P^q^*. These two quantities evolve according to a Markov chain with transitions corresponding to processes such as filament branching, capping, severing, splitting and uncapping (corresponding rates indicated as *γ*(*t*) in Fig. 1B), mediated by actin-binding proteins such as Arp2/3, CapZ and cofilin (Blanchoin et al., 2000; Borovac et al., 2018; Bonilla-Quintana et al., 2020). Due to crosslinking proteins, actin filaments can be converted to a relatively slower, stable actin pool (*S* in Fig. 1B&D), modelled as a low-pass filtered version of the dynamic pool. To model LTP-induced changes in dynamic and stable actin, we adjust the rates of the different processes according to the relative concentration changes of the associated actin-binding proteins (Bosch et al., 2014, see Methods 4.2). The combined action of these two pools generates an expansive force (*F_act_* in Fig. 1D, see Methods 4) that drives spine enlargement under LTP. The spine membrane generates a counterforce (*F_mem_* in Fig. 1D) as it resists the changes of the spine volume, surface area, and local curvature (Helfrich, 1973; Guckenberger and Gekle, 2017).

To extend this model by the geometry-gated PSD dynamics, we need to determine whether the membrane is tightly wrapped around the PSD perimeter and blocks growth, or whether it gives the PSD sufficient space to grow. For this, we need measure of the local membrane-curvature at each point *i* of the PSD-periphery. For computational efficiency, we utilize the respective curvature-induced membrane-force component *f_C_*(*i*), which is proportional to the local mean curvature. Growth is blocked at any point of the PSD periphery, if there is a large curvature bending the membrane towards the PSD (see Fig. 1E, condition 3). As our PSD is parallel to the x-y-plane, this means that at point *i*, the z-component of the curvature force *f_C,z_(i)* is negative (inward directed). Also the force in the PSD-growth direction *f_C,n_(i)*, which corresponds to the normal vector of the PSD-perimeter in the x-y-plane *n_xy_*(*i*) (see Fig. 1C), needs to exceed a certain threshold *θ* to block the insertion of new scaffolds at the PSD. If this force is smaller than the threshold (condition 1 in Fig. 1E), or the membrane bends upward and *f_C,z_ >* 0 (condition 2 in Fig. 1E), growth is allowed.

Additionaly, PRPs must be available allow for the outgrowth of the PSD. Thus, the above geometrical conditions is coupled with the availability of PRPs to achieve protein capture as summarized in Table 1.

**Table 1:** Protein capture gated by membrane geometry and PRP availability.

| Membrane at the PSD periphery | PRPs | No PRPs |
| --- | --- | --- |
| Constrained: $f_{C,n}(i) \geq \theta$ and $f_{C,z}(i) < 0$ | **✗** | **✗** |
| Permissive: $f_{C,n}(i) < \theta$ or $f_{C,z}(i) \geq 0$ | **✓** | **✗** |

For the PRP availability due to either local or global protein synthesis (Bramham, 2008; Bradshaw et al., 2003), we use a time-dependent step function.

Taken together, protein capture in our model is represented by PSD growth, and is governed by the interplay between protein availability and membrane curvature, which here acts as the structural interpretation of the synaptic tag.

### Actin-spine geometry-based synaptic tag captures PRPs by facilitating PSD growth

In the following, we examine the behaviour of this tagging and capture mechanism under LTP stimulation. To this end, we simulated the model under LTP conditions and observed the resulting changes in dynamic and stable actin, curvature-induced membrane counterforce components, spine volume and PSD area (Fig. 2). Prior to LTP, the model equilibrates to reach a basal/control state (time until *t* = 0.0 h in Fig. 2). Upon LTP, the stimulus (Fig. 2A) induces a rapid increase in dynamic actin, reflected in the total number of barbed ends *B* across all active foci in the spine (*B* in Fig. 2B). This is transient and decays quickly, while a slower increase in stable actin also builds up in parallel which decays much more slowly (*S* in Fig. 2B). This rapid and slower responses of actin pools work together with the spine membrane to bring about the increase in spine volume (Fig. 2E, blue) observed within the first 60 minutes after LTP.

**Figure 2:**
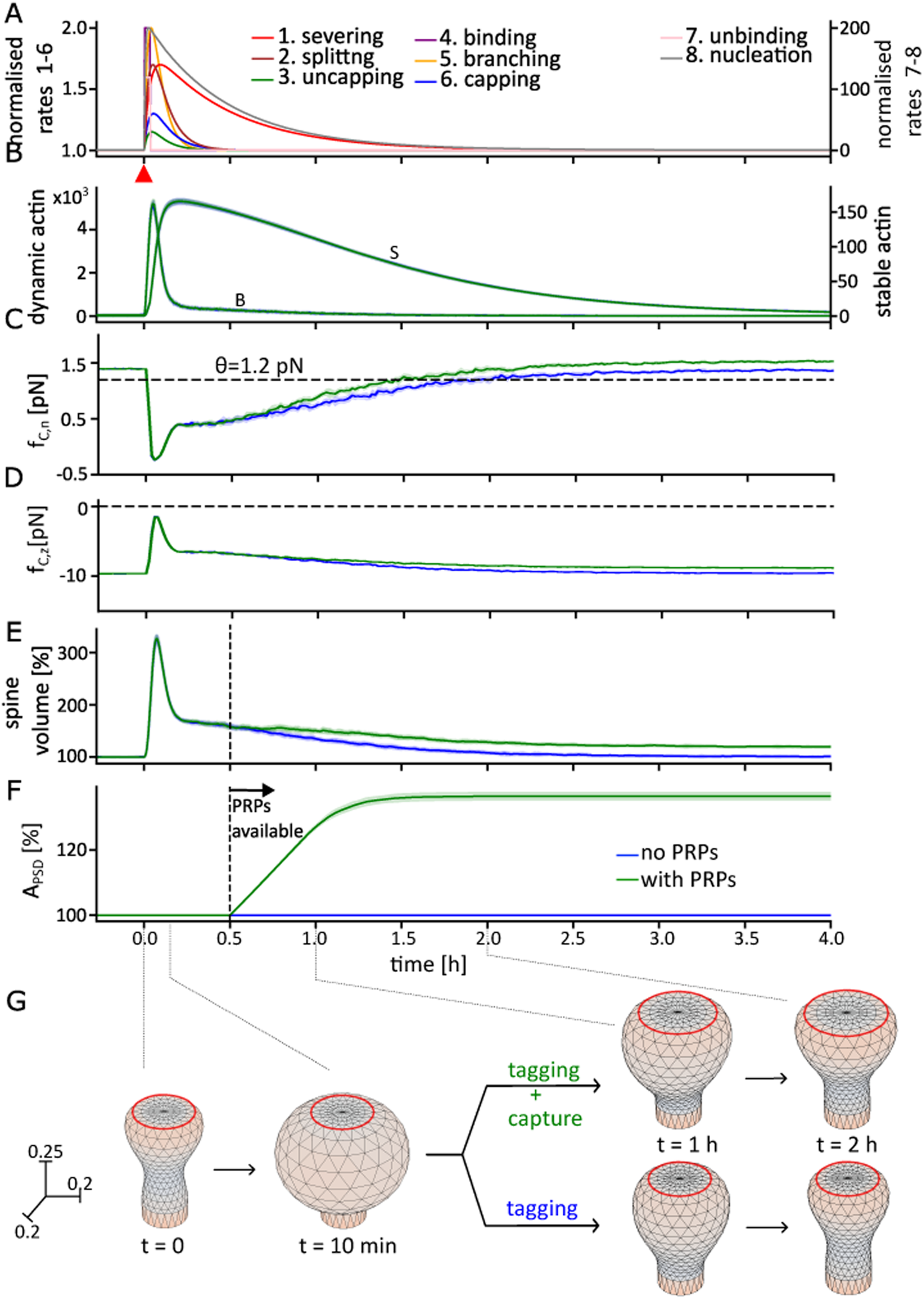
Synaptic tagging and capture in LTP. (A) Time-dependent rates of the different ABP-related processes following an LTP-inducing stimulus (red triangle at *t* = 0h). (B) Time course of dynamic actin, measured as the total number of barbed ends from all active foci *B*, and the size of the slower stable actin-pool *S*. Curves and shaded areas in panels B - F depict mean and standard deviations of the quantities over 20 trials. Blue and green curves indicate simulations without and with PRP availability, respectively.(C) In-plane component of the curvature induced membrane force averaged over all PSD periphery vertices. (D) Same for the *z* component. (E) Spine volume exhibits a fast overshoot and a slower decay, which is converted to lasting volume increase when PRPs become available (vertical dashed line at *t* = 0.5 h). (F) Area of the PSD (relative to its initial value) starts to increase when PRPs become available. (G) triangular meshes of the spines at different time instants along two simulations. The red ring at the top of each snapshot represents the PSD periphery, which enlarges upon PRP capture.

This spine enlargement produces a shift in both the in-plane resistance to lateral expansion of the PSD *f_C,n_* and the out-of-plane curvature component at the PSD periphery *f_C,z_*. The out-of-plane force transiently increases towards more positive values and then decreases (Fig. 2D), while the in-plane force transiently decreases and gradually recovers as the membrane relaxes (Fig. 2C). During this transient decrease, the spine fulfils the condition for protein capture (condition 2 in Fig. 1E). Upon PRP availability (here, from *t* = 0.5 h for the green curves), the PSD starts to expand and its area increases (Fig. 2F). This process continues until the curvature force-components fail to fulfil the defined condition and the PSD expansion ceases. Due to the PSD expansion, also the spine volume converges to a higher value (Fig. 2E, green) as compared to the condition without PRP availability (Fig. 2E, blue).

Notably, although PRP availability is maintained throughout the remainder of the simulation, it results in PSD growth only as long as the geometrical conditions are satisfied, which is not the case in the basal state or long after LTP. This is consistent with the STC principle that PRPs can only be captured at tagged synapses. Together, these results demonstrate that local spine geometry information around the PSD could act as a tagging mechanism facilitating protein capture, thereby converting a transient actinspine-geometry-based state into a persistent state resembling l-LTP.

### Timing of PRP availability decides the degree of synaptic consolidation

In the following, we further characterise our model by taking into account the effects of input timing. Experiments using a weak-before-strong protocol showed that synapses which are tagged by a weak stimulation (WTET) but do not elicit PRP synthesis, can still exhibit l-LTP if PRPs later become available due to another, strongly stimulated synaptic pathway (using strong tetanisation, STET) (Frey and Morris, 1997). The time interval between the two stimulations decides the extent of potentiation of the WTET pathway, indicating a continuous decay of the synaptic tag. To test whether our proposed geometry-based tagging and capture mechanism aligns with the findings of this protocol, we investigated varying time delays between the LTP stimulus and the onset of PRP availability. Here we assume that our LTP stimulus results from applying a weak stimulation and only sets the geometric tag, and the PRP availability signals are assumed to be the outcome of STET applied elsewhere with a certain delay. The growth condition for the spine geometry is fulfilled shortly after LTP onset (Fig. 1C). Thus, when the PRPs are available earlier, the PSD growth lasts longer (Fig. 3D, purple curves) until the growth condition is not met anymore. The longer the PRP availability is delayed, the shorter the interval where the growth condition is met, leading to less PSD growth. This correspondingly lowers the magnitude of increase. The resulting spine volume reflects this (Fig. 3A). Maximal PSD growth and consolidation is achieved if the PRPs are immediately available (compare Negri et al. (2025)) after the LTP stimulus. The PSD growth drops to zero if the PRP availability starts after the tag has decayed completely (Fig. 3E-F). Spine volume shows the same dependence in its post-stimulation steady state: while all conditions exhibit a transient increase following LTP (Fig. 3A), the enlarged state is sustained only when PSD growth occurs; otherwise, the volume relaxes towards baseline (Fig. 3B-C).

**Figure 3:**
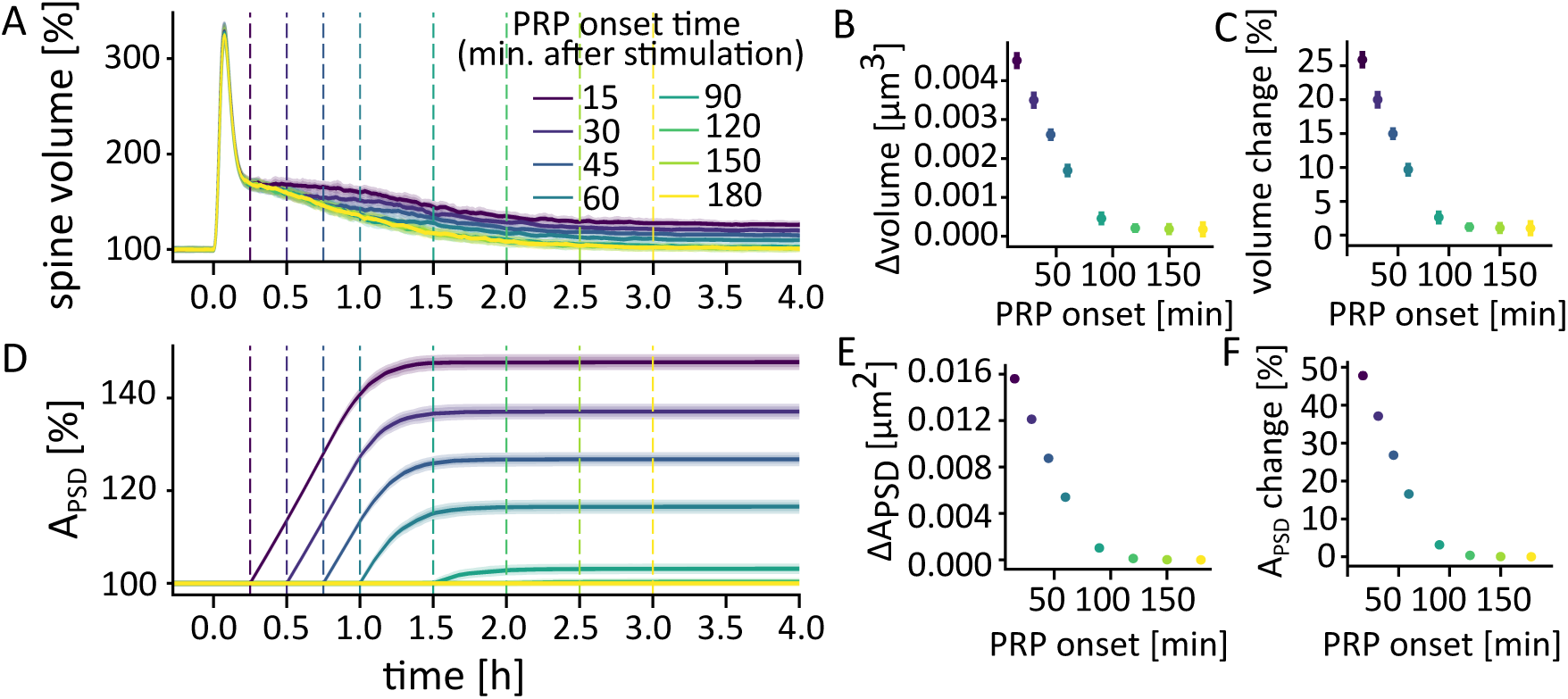
Different delays in PRP availability and the resulting changes (results are averaged over 20 trials). (A) Spine volume changes for different PRP onset times in minutes (see legend, also dotted lines) after applying the LTP stimulus. (B) Absolute changes in spine volume. (C) Percentage changes in spine volume, calculated as (steady state post-LTP volume - basal volume)/basal volume. (D) PSD area for different PRP onset times. (E) Absolute changes in PSD area. (F) Percentage changes in PSD area, calculated as (steady state post-LTP area - basal area)/basal area

These results show that effective PRP capture is constrained by the lifetime of the actin-based geometrical tag, thereby defining an operating regime in which protein availability must overlap with the permissive geometrical state. The duration of this overlap determines the degree of synaptic consolidation.

### Geometry-dependent timing of spine volume and PSD growth

So far we have investigated the influence of input timing on the final LTP outcome. However, experiments show that the outcome of plasticity also depends on the initial size of the spine, with small spines being more prone to modifications, and bigger, mushroom spines more resistant to further potentiation (Helm et al., 2021; Kasai, 2023). In our model, different initial spine and PSD sizes give rise to different initial curvatures around the PSD and also different time-intervals during which the geometric conditions for growth are met. We thus expect that size-dependent plasticity arises naturally from the geometrical consolidation mechanism we proposed.

To test this, we first simulated the geometry dynamics for different fixed PSD sizes in the absence of PRP-dependent PSD remodelling (Fig. 4A-D). For the steady state, this revealed that the bigger the PSD, the higher the in-plane membrane counterforce component *f_C,n_* at the PSD periphery (Fig. 4B). Also, the component *f_C,z_*shows a saturation pattern as we move to bigger PSD sizes, reflecting reduced upward membrane bending around larger PSDs (Fig. 4D). Upon LTP (*t* = 0.0h in Fig. 4A & C), all spines show positive change in *f_C,z_* (Fig. 4B), while *f_C,n_* exhibits a dip and slow recovery to the initial value. Also, the basal value of *f_C,n_* is smaller for smaller PSD sizes such that the geometric growth condition is fulfilled longer. This indicates that smaller PSDs are susceptible to growth for longer durations.

**Figure 4:**
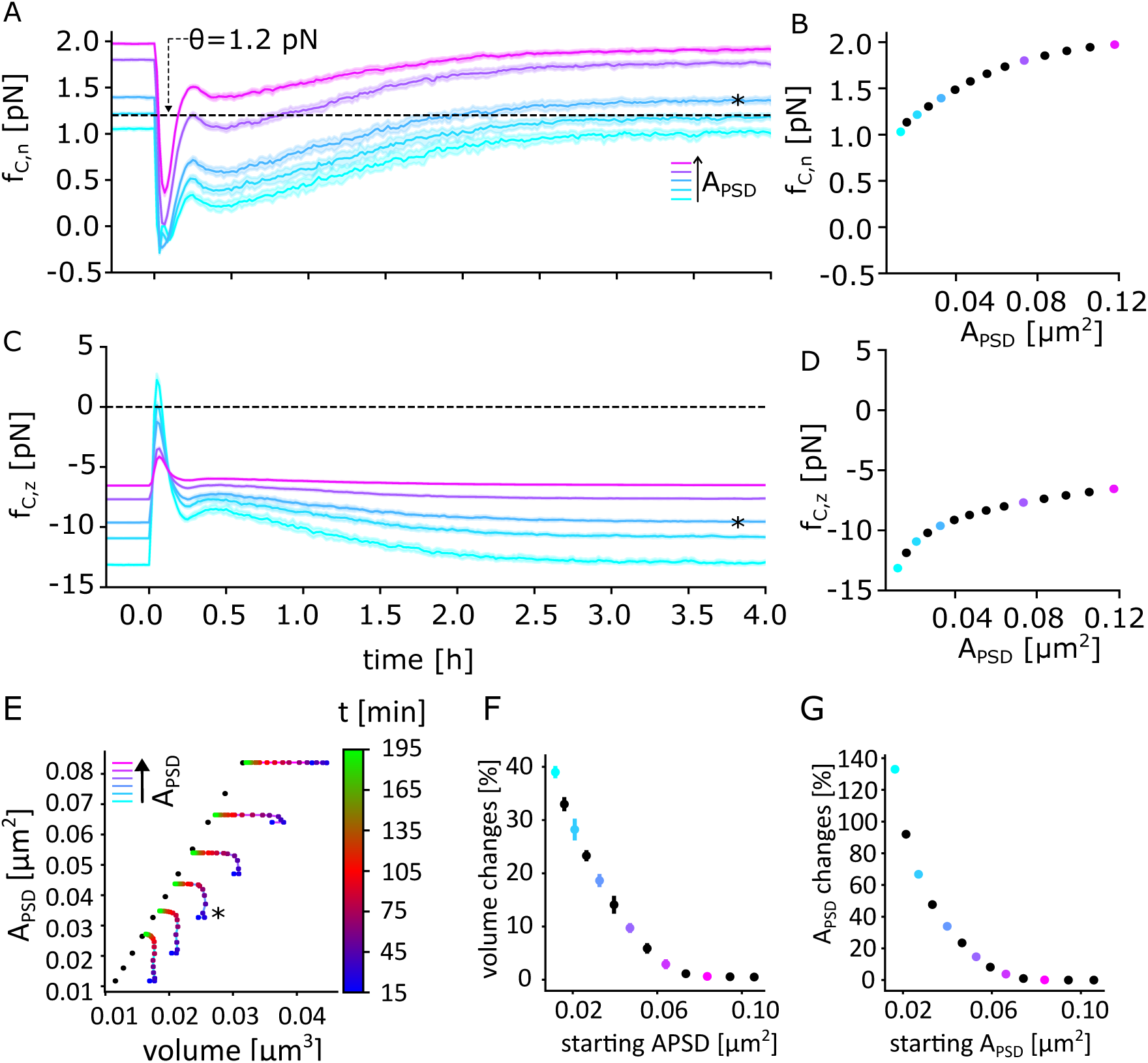
Influence of PSD size on membrane forces and plasticity outcome. (A) Time-course of the in-plane projection of the curvature-induced counterforce for different PSD sizes under LTP-conditions but without PSD dynamic. Curves and shades depict mean and std for 20 simulations, respectively. Shortly after LTP-onset, a sharp decrease, followed by a rebound and a slow relaxation can be observed. Dashed line depicts the threshold for gating PSD growth (*θ* = 1.2pN). Curves of smaller spines stay below the threshold longer than larger spines. Asterisk marks the PSD size used in previous figures. (B) Steady state value of the in-plane force (without LTP). (C, D) Same for the z-component of the curvature-induced counterforce. (E) LTP-induced trajectories in the spine volume-PSD space for spines with different initial PSD sizes and PSD dynamic enabled. Dot colour indicates time after stimulation (see colour-bar) while the connecting line colour indicates initial PSD size. Black dots indicate basal values for different PSD sizes and exhibit the experimentally observed correlation. This correlation is transiently shifted towards larger volumes early after LTP. (F) Relative spine volume changes for different initial PSD sizes. Larger spines show relatively less LTP. (G) Same for relative PSD size changes.

Based on these observations, we expect to see progressively smaller changes in spine volume and PSD for bigger PSD sizes under PRP availability. To test this, we simulated LTP for all the PSD sizes with PRP availability. Indeed, we observed that the relative changes in spine volume and PSD area reduce in magnitude with increasing initial PSD sizes (Fig. 4E-G). Thus, our model predicts a saturation of LTP for very large spines, while smaller spines are very plastic.

As our model exhibits the experimentally observed correlation between PSD size and spine size in basal state (Bonilla-Quintana et al., 2020), we wondered how this relation evolves after LTP. In experiments, the PSD size and spine volume correlation is briefly broken (or shifted) after LTP before it is eventually re-established (Sun et al., 2021; Bosch et al., 2014; Meyer et al., 2014). We tested whether our model can reproduce this phenomenon, and assessed the PSD-spine volume trajectory starting from shortly before the PRPs become available until three hours later (colour bar in Fig. 4E). Soon after stimulation, we see that the correlation of the basal values (black dots) is broken in favour of larger volumes (rightward shift, compare Bosch et al., 2014; Sun et al., 2021). The trajectories return to the basal correlation line, but settle at a new steady state with increased spine and PSD size. Here again, higher magnitudes of changes are observed for smaller initial PSD sizes (Fig. 4F). The spine enlargement is matched by the PSD enlargement over time for this re-establishment of correlation (Fig. 4E, G). The same relation is observed between the relative increments in volume and PSD as well (Supp. Fig. S2A).

Taken together, these results imply that the geometrical tagging mechanism also aligns with the size-dependent synaptic plasticity outcomes, because bigger spines interact weakly with the PRP availability as their membrane curvature fulfils the conditions for PSD growth for shorter time intervals.

## 3 Discussion

In this study, we proposed a PRP capture mechanism based on the assumption that capture-related PSD growth can be gated by the spine geometry, in particular the curvature of the spine membrane around the PSD. We demonstrated that a model based on this mechanism indeed yields PSD growth after LTP. We further analysed how this growth depends on the timing of protein availability as well as on the initial size of the spine, obtaining predictions which are qualitatively comparable to experiments. Taken together, these results support a view of the synaptic tag not as a single molecular entity, but as a transient structural and mechanical state of the synapse encoded by actin dynamics and spine geometry (compare Pinho et al., 2020; Okuda et al., 2021; Redondo and Morris, 2011).

A summary of the proposed geometry-based STC mechanism is shown in Figure 5. Coarsely speaking, the state of the spine is determined by interactions of actin dynamics, spine geometry and the size of the PSD (blue boxes). Under basal actin dynamics, the PSD determines the spine size (Fig. 5, iv and vi, compare Bonilla-Quintana et al., 2020). When LTP stimulation is applied, the dependence on PSD size is relatively weakened and the spine enlarges (Matsuzaki et al., 2004; Meyer et al., 2014; Sun et al., 2021) through increased remodelling of dynamic actin (Fig. 5, ii; compare Bosch et al., 2014). This results in a transient increase in stable actin and altered spine geometry on the timescale of hours (Thomas et al., 2025), which marks the tagged state. Besides increased spine volumes, this also entails curvature variations of the spine membrane around the PSD. If the spine membrane is upward bending or the projection of the membrane counterforce into the PSD-plane is small enough, the PSD extends laterally upon PRP availability (Fig. 5, iii; compare Frey and Morris, 1997, 1998). Thus, this geometric condition can be viewed as the biophysical implementation of the tag and PSD growth only occurs if this tag is present. Once actin dynamics returns to its basal state, the increased PSD size also entails a long-lasting increase in spine volume (Fig. 5, iv). Without available PRPs within the lifetime of the tag, however, the PSD does not grow (Fig. 5, v) and the spine goes back to the pre-stimulation state (Fig. 5, vi; compare Thomas et al., 2025). A testable prediction of this mechanism would be that long-term potentiation or spine enlargement should be correlated with membrane curvature around the PSD shortly after LTP. To measure this, longitudinal super-resolution experiments (Clavet-Fournier et al., 2024; Sun et al., 2021) would be needed to provide enough spatial resolution and glutamate uncaging (Ellis-Davies, 2019) could provide sufficient temporal resolution of the timing of the stimulus.

**Figure 5:**
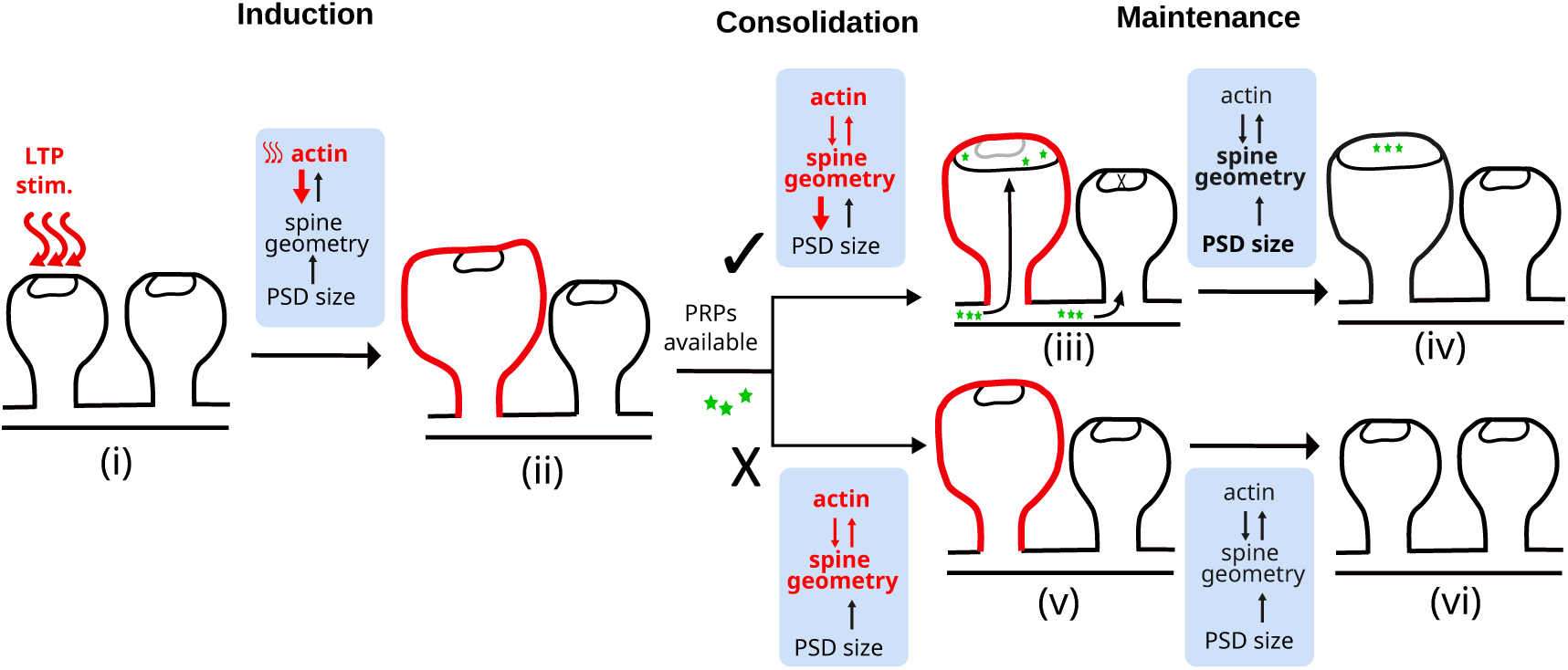
Conceptual summary of the geometry-based synaptic tagging and capture.

In our model – similar as in experiments (Matsuzaki et al., 2004; Meyer et al., 2014; Helm et al., 2021) – long term changes of the PSD are dependent on the initial spine and PSD size. This can be explained with mechanics: for larger PSDs, the curvature component of the membrane counterforce is always strong as the membrane must curve more to connect the neck with the rim of the PSD. Hence, large spines do not satisfy the condition for further PSD growth. Within the STC framework, the deformation of larger spines can no longer function as an effective tag for PRP capture. This suggests that spine geometry, and by extension its morphology, may encode a biophysical boundary between plasticity and stability — a structural correlate of memory protection or write-protection mechanism for larger spines (Kasai, 2023). An interesting extension of our model, which could impact these findings is neck plasticity(Tønnesen et al., 2014; Gupta and O’Donnell, 2023; Araya et al., 2014), which we so far neglected due to the relatively stable, ringlike actin cytoskeleton in the neck (Bär et al., 2016). A plastic neck, however, would change the lower anchoring points of the spine head membrane and therefore influence the curvature landscape and thus plasticity outcome in our model.

Varying the timing between the tag setting and the PRP onset can result in different plasticity outcomes (Frey and Morris, 1998). Here we presented a mechanical interpretation for these differences (see Fig. 2C, D): without PSD-growth, spines gradually decay back to their previous state, thereby adapting the curvature around the membrane until growth is eventually blocked. The outcome of plasticity is determined by how much time is left until this time-point at the onset of PRPs, and is therefore continuously decreasing over time. The timescales of the geometric tag-PRP interactions yielding visible increases are in the range below two hours. Here, we analysed this time-window for a single, initial spine size, while it can be assumed that the synapse population measured in experiments consists of a broad spectrum of spine sizes. As the initial size also determines the outcome of plasticity, one would have to consider a population of spines in order to reproduce the experiments faithfully. Indeed additional simulations showed that smaller spines have a longer tagging interval that exceeds two hours, whereas large spines have shorter tagging intervals (see Supplementary Figure S1). Moreover, there could be further factors which help extend this window. For instance, the mechanical interactions with the presynaptic terminal (Meyer et al., 2014; Fisher-Lavie and Ziv, 2013), extracellular matrix (Dankovich and Rizzoli, 2022), and astrocytes/glia (Freche et al., 2011) would impact the geometry dynamics described here and potentially aid in extending the timescale for viable tag-PRP interactions.

Finally, our study focused on LTP, while the analogue mechanism for LTD (Szabó et al., 2016) remains an open question. Here, we would speculate that an LTD-induced downregulation and depolymerisation of actin could lead to the experimentally observed spine shrinkage (Zhou et al., 2004), which would put even stronger forces onto the PSD- periphery. The LTD-related PRPs would then have to allow for PSD shrinkage, e.g., by dissolving the mesh of PSD-scaffolds (Camus et al., 2025). Similar as here, also PSD shrinkage could then be modelled by a force threshold plus PRP availability. Along this line, an abstracted version of the here presented model (Negri et al., 2025) has been applied to explain both LTP- and LTD-related tagging and capture experiments.

Yet, also the picture of a two-dimensional, layered PSD may be too simple. Recent studies report that applying a plasticity-inducing stimulus reorganized PSD molecules from a single cluster into multiple clusters (Christensen et al., 2022; Zeng et al., 2018), which complies with the notion of perforated and fragmented PSDs post-stimulation (Stewart et al., 2005). To study the impact of this, a more fine grained PSD-dynamics, which captures these effects on a molecular scale (Safari et al., 2025), could be integrated with our membrane and actin dynamics in the future, which would also allow to test our assumptions on the Brownian ratchet mechanism.

In conclusion, our presented work proposes a geometrical basis for synaptic tagging and capture, which could also be used as basis to analyse pathological conditions where aberrant dendritic spine morphology and actin dynamics results in neuropsychiatric disorders such as autism spectrum disorder, schizophrenia, etc. (Borovac et al., 2018). Decoding how the geometrical landscape of spines is tuned or disrupted and how this impacts synaptic plasticity and learning could open new directions for understanding not only the cellular basis of cognition but also the origins of its failure in disease.

## 4 Methods

We base our work on a published implementation of the LTP-induced dynamics of actin and the spine membrane Thomas et al. (2025). Except for the PSD-remodelling and PRP availability, all equations and parameters remain the same. However, for the sake of completeness, we here repeat the major model equations and assumptions, but skip details on the numerical implementation.

### Model of the spine membrane

Our membrane model is a movable 3D triangular mesh(Thomas et al., 2025). It consists of 382 vertices arranged to resemble a dendritic spine with a fixed neck region. Starting with a sphere of radius 0.25*µ*m divided into 20 longitudinal and latitudinal sections, the top four sections are flattened to form the PSD disc and the bottom three are made into a cylinder to form the neck. Each vertex that constitutes the structure is acted upon by actin-generated forces from within and counterforces from the membrane. The imbalance between these two decide the magnitude and direction of motion for the vertices in the spine head (excluding PSD) and hence the overall changes in the spine morphology. The vertices of the neck region remain fixed, since the neck is reported to be composed of more stable actin rings (Bär et al., 2016; Korobova and Svitkina, 2010).

The PSD is represented by the vertices in the flattened section and we assume it to remain a planar structure that only remodels or expands at its periphery and inside the plane due to the below described dynamics. This abstraction is based on experimental evidence of the layered structure with a relatively stable horizontal mesh of scaffolding proteins (Blanpied et al., 2008; Chen et al., 2008; Hayashi et al., 2009).

### Actin dynamics

#### Dynamic pool of actin

We assume that the spine contains two pools of actin: dynamic and stable (Honkura et al., 2008). The dynamic actin pool is assumed to consist of a time-varying number of actin polymerisation foci Frost et al. (2010).

New foci are nucleated depending on the nucleation rate *γ_nucl_*(*t*). We choose their locations probabilistically as follows (i) a set of candidate nucleation points is generated by forming an inner point cloud (generated by scaling down the vertex coordinates of the spine head excluding the PSD and neck by a factor of 0.8); (ii) one of them is chosen for newly nucleated focus with a probability that exponentially decays with their distance from the PSD

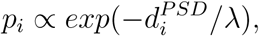

where 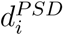 is the Euclidean distance of candidate point i from the centre of the PSD and λ a scale parameter (Table 4).

The state of each focus *q* is described by its number of uncapped barbed ends *B^q^* and uncapped pointed ends *P^q^* with random initial values(Bonilla-Quintana et al., 2020; Thomas et al., 2025). At each time step, each focus undergoes one or more of the different actin-associated processes with an associated rate *γ.*. These rates are multiplied with *dt* = 0.05, to obtain probabilities for each associated process in each time step. The processes update the number of barbed and pointed ends as follows:

- *Branching (B^q^ → B^q^*+ 1): Attachment of ARP2/3 to an existing filament can nucleate a new branch, creating an additional barbed end. The rate of branching is given by

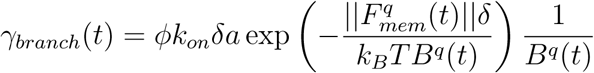 where ϕ is a proportionality constant, k_on_ is the G-actin assembly rate constant, a is the available actin concentration, 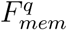 is the counteracting membrane force at the centre of focus *q*, δ is gained filament length by adding a single G-actin molecule, *k_B_T* is the thermal energy, and *B^q^*(*t*) the number of barbed ends at time t (Table 4).
- *Capping (B^q^ → B^q^ −* 1): Binding of a capping protein (e.g., CapZ) to a barbed end halts its polymerisation, effectively removing that barbed end from active growth.
- *Severing (B^q^ → B^q^ −*1*, P^q^→ P^q^ −*1): Cofilin binding can sever a filament, followed by complete depolymerisation of the severed segment.
- *Splitting (B^q^ → B^q^* + 1*, P^q^ → P^q^* + 1): Under certain conditions, cofilin-mediated severing can also generate a new active filament, increasing both barbed and pointed ends.
- *Uncapping (P^q^ → P^q^* + 1): When ARP2/3 dissociates from a capped pointed end, it produces an uncapped pointed end that is free to depolymerise.

A focus terminates and is removed from the simulation when the condition B^q^ = 0 is reached.

#### Stable pool of actin

Stable actin is formed by the binding of crosslinking proteins to dynamic filaments (Honkura et al., 2008; Shaw et al., 2021). We assume this process happens at a certain binding rate *γ_bind_*(*t*). Also, it decays upon the unbinding of crosslinkers, through an unbinding process with rate *γ_unbind_*(*t*). We implement this without explicitly feeding it back into the dynamic pool (Table 3). The stable actin dynamics is hence modelled as

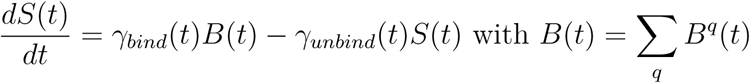

where *B* here is the total number of barbed ends summed over all active foci at the given time *t*, and *S* the amount of stable actin.

#### Actin-generated force

The actin-generated force *F_act_* is assumed to stem from the retrograde movement of filaments that polymerise at the chosen locations of nucleation. At each vertex, contributions from all foci, which grow into its direction are summed. Hereby, the force from a certain focus *q* onto a vertex *i* is proportional to its barbed ends and is given by

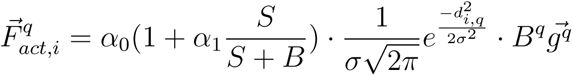

where *B^q^* is its total number of barbed ends, and 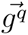 is the focus growth direction, which we have chosen for simplicity, to be along the line though the nucleation point and origin. The Gaussian term distributes the force from each actin focus with a spread *σ* on the mesh, so a single focus can exert force on multiple vertices. Hereby, *d_i,q_* is the orthogonal distance of vertex *i* w.r.t the growth direction of the focus *q*. Stable actin influences the spine geometry by resisting the retrograde movement of filaments back towards the center of the spine. To model this effect, we use the prefactor 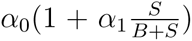 which introduces the additional contribution from the stable actin fraction 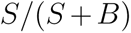 scaled by the parameter *α*_1_ (Table 4).

### 4.1 Membrane dynamics

The displacement of vertex *i* with respect to time is

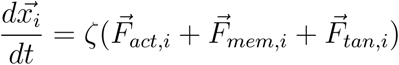

where *ζ* is the speed of update (Table 4). The membrane counterforce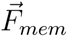 is calculated following the Canham-Helfrich free energy formalism for thin lipid membranes (Helfrich, 1973; Canham, 1970; Thomas et al., 2025; Bonilla-Quintana et al., 2020), as the spatial gradient of bending energy *E*, which consists of contribution from the changes in volume, area and curvature.

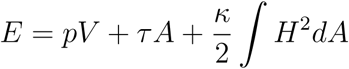

Here, *p* is the pressure difference between intracellular and extracellular sides, *V* the spine head volume, *τ* the surface tension, A the surface area, *κ* the bending modulus, and *H* the mean curvature. Note that *p* and *τ* are considered to be constant as the spine is open to the dendrite. The membrane counterforce at each vertex *i* then becomes

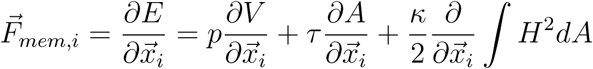

where 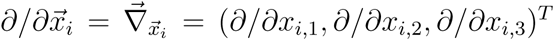 is the derivative with respect to the individual coordinates of vertex i.

As the focus of the present paper is membrane curvature, we here only sketch the curvature contribution. The mean curvature integral in the above equation can be discretised/numerically approximated as

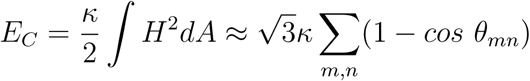

where *θ_mn_* is the dihedral angle between two adjacent triangles *m* and *n* (Guckenberger and Gekle, 2017). This in turn, can be written in terms of the face normals 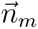 and 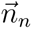 as

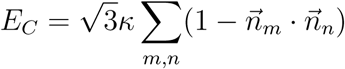

Each vertex *i* participates in several adjacent triangle pairs *m, n* that form a local triangle constellation around it. The total curvature energy *E_C_*is the sum of the contributions from all such neighbouring face pairs. The curvature contribution to the membrane counterforce on vertex *i* is then given by the derivative of the above approximation to the curvature energy:

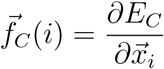

In addition to these physical force components, we include a geometric constraint term 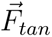, redistributing vertex displacements tangentially to the spine surface to prevent excessive local distortion during updates, keeping the mesh triangles more or less equilateral.

### 4.2 LTP stimulus

Under basal conditions, the rates associated with each ABP *γ_branch_*(*t*), *γ_cap_*(*t*), *γ_uncap_*(*t*), *γ_sever_*(*t*), and *γ_split_*(*t*), as well as the nucleation rate *γ_nucl_*(*t*) are kept at a constant value. Under LTP, it is reported that ABPs undergo variations in their concentrations in spines (Bosch et al., 2014). Mathematically, their time courses resemble a double exponential function that has a rising and falling edge with distinct time constants. Hence, we use such a description to model the rates associated with ABPs as follows:

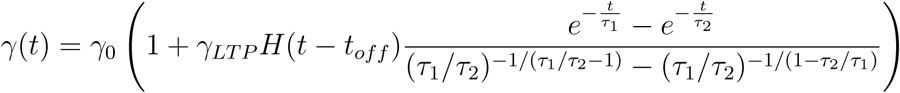

where *γ*_0_ is the rate under basal condition, *γ_LTP_* the change relative to the basal rate, *τ*_1_ the rising time constant, and *τ*_2_ the falling time constant, *H* the Heaviside function, and *t_off_* the time of LTP stimulation. The parameters used to model each rate are given in Table 2.

**Table 2:**
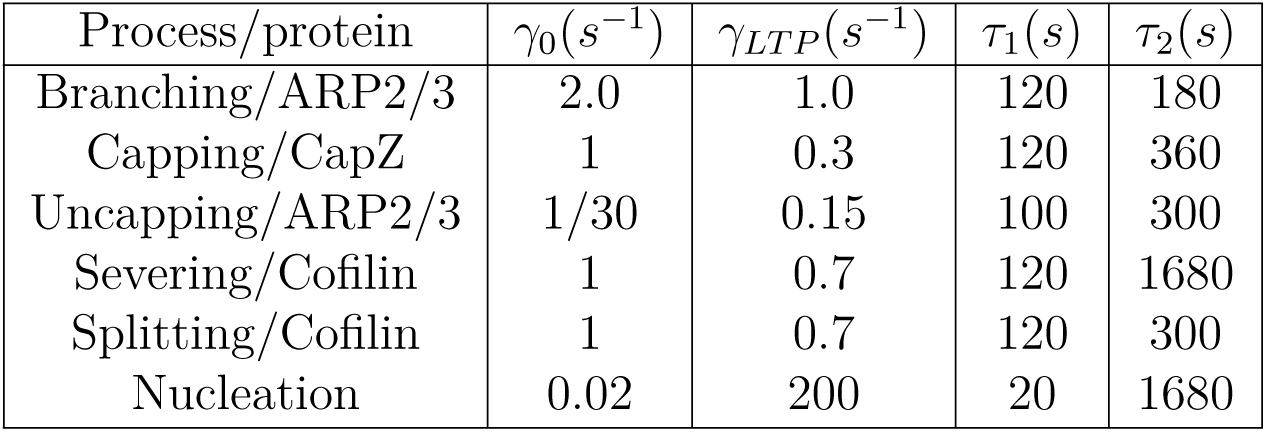
Time-dependent model parameters varied by LTP-inducing stimulation taken from Thomas et al. (2025)

| Process/protein | $\gamma_0(s^{-1})$ | $\gamma_{LTP}(s^{-1})$ | $\tau_1(s)$ | $\tau_2(s)$ |
| --- | --- | --- | --- | --- |
| Branching/ARP2/3 | 2.0 | 1.0 | 120 | 180 |
| Capping/CapZ | 1 | 0.3 | 120 | 360 |
| Uncapping/ARP2/3 | 1/30 | 0.15 | 100 | 300 |
| Severing/Cofilin | 1 | 0.7 | 120 | 1680 |
| Splitting/Cofilin | 1 | 0.7 | 120 | 300 |
| Nucleation | 0.02 | 200 | 20 | 1680 |

Additionally, we model the crosslinker binding and unbinding rates *γ_bind_* (*t*) and *γ_unbind_* (*t*) with a double-step function.

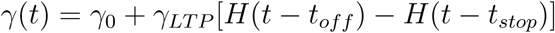

where *t_stop_* the time when the crosslinker rates get back to the baseline values (2 minutes post-stimulation). The parameters used to model these rates are given in Table 3.

**Table 3:**
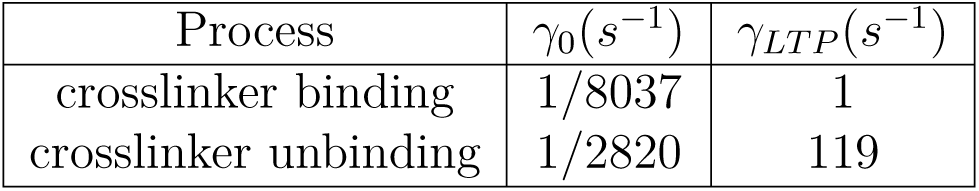
Time-dependent model parameters varied through LTP for crosslinkers taken from Thomas et al. (2025)

**Table 4:** Time-independent model parameters. All except *ζ_PSD_* taken from Bonilla-Quintana et al. (2020); Thomas et al. (2025)

| Name | Quantity | Value | Unit |
| --- | --- | --- | --- |
| proportionality constant | $\phi$ | 75 | $\mu m^{-1}$ |
| G-actin assembly rate constant | $k_{on}$ | 11.6 | $\mu M^{-1} s^{-1}$ |
| length of G-actin | $\delta$ | 0.0022 | $\mu m$ |
| thermal energy | $k_B T$ | 0.0041 | $pN \mu m$ |
| nucleation distance parameter | $\lambda$ | 0.1 | $\mu m$ |
| actin force amplitude | $\alpha_0$ | 0.5 | $fN$ |
| stable pool influence on force | $\alpha_1$ | 10 | - |
| spatial spread of each focus | $\sigma$ | 0.05 | $\mu m$ |
| bending modulus | $\kappa$ | 0.18 | $pN \mu m$ |
| surface tension | $\tau$ | 15 | $pN \mu m^{-1}$ |
| diff. between internal and external pressure | $p$ | 85.7143 | $pN \mu m^{-2}$ |
| speed of mesh movement | $\zeta$ | 0.002 | $\mu m^2 s^{-1} pN^{-1}$ |
| speed of PSD movement | $\zeta_{PSD}$ | 0.0000075 | $\mu m^2 s^{-1} pN^{-1}$ |

### 4.3 PSD dynamics and PRP availability

PSD remodelling is constrained by curvature-dependent conditions described above. Specifically, PSD expansion is permitted if (i) the membrane at the PSD periphery exhibits upward bending or (ii) the in-plane component of the membrane counterforce remains below a threshold *θ*. These conditions are implemented to update the position of each PSD vertex *i* as follows.

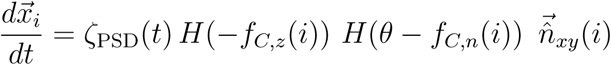

where *f_C,z_* (*i*) is the z component of the curvature induced force 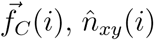 the in-plane normal vector to the PSD periphery, and *f_C,n_*(*i*) the force component in this direction.

We include the PRP availability in the growth speed ζ_*PSD*_ of the PSD periphery, resulting in

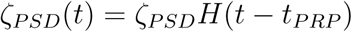

where *t_PRP_* the time that the PRP availability starts. Thus, when the PRPs are available and the local geometry around the PSD conducive for their addition, capture would take place.

### 4.4 Software implementation and code availability

We used Python and Numpy for simulating the model and analysing the results. Close to real-time simulation times were possible on a consumer PC for individual simulations. Multiple simulations were run in parallel on the supercomputer cluster of the Gesellschaft für wissenschaftliche Datenverarbeitung mbH Göttingen (GWDG).

## Acknowledgments

This work was funded by the German Science Foundation under CRC1286 “Quantitative Synaptology”, project C03. We would like to thank Stefan Klumpp, Jannik Luboeinski, Silvio Rizzoli, Christian Tetzlaff and Florentin Wörgötter for fruitful discussions on the project and Lennart Jahn for helping with the operation of the compute cluster.

