## Supplementary Information for "Geometry-based dynamics of the postsynaptic density explain protein capture by an actin-spine-geometry-dependent synaptic tag"

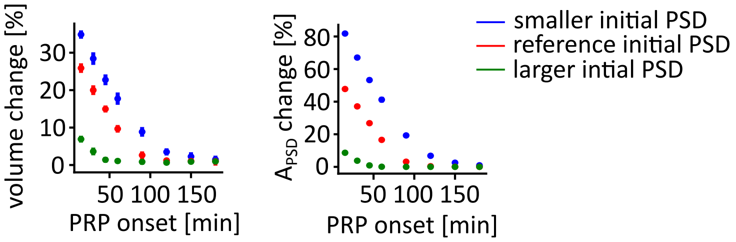


Figure S1: *Initial spine and PSD size and PRP onset time jointly determine the plasticity outcome* . Relative change of spine and PSD size under varying PSD onset times for the spine size in Fig. 3 (red) as well as larger (green, 150% scaling) and smaller (blue, 80% scaling) initial spine and PSD size

| scaling factor from reference | area in $\mu m^2$ |
|---|---|
| 0.6 | 0.012 |
| 0.7 | 0.016 |
| 0.8 (small spine) | 0.021 |
| 0.9 | 0.026 |
| 1.0 (reference size) | 0.033 |
| 1.1 | 0.040 |
| 1.2 | 0.047 |
| 1.3 | 0.055 |
| 1.4 | 0.064 |
| 1.5 (large spine) | 0.074 |
| 1.6 | 0.084 |
| 1.7 | 0.094 |
| 1.8 | 0.110 |
| 1.9 | 0.118 |

Table S1: Different initial PSD areas used in the simulations

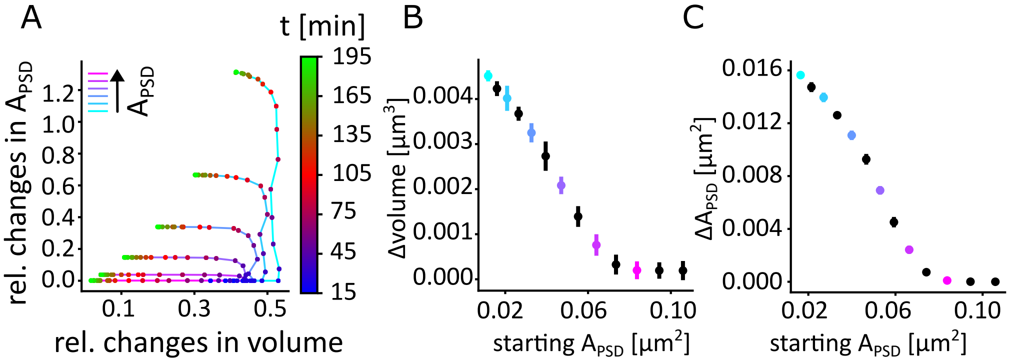


Figure S2: *Initial spine and PSD size determines the extent of potentiation.* (A) Relative change of spine volume and PSD area for different initial PSD areas. Color of the points indicates the time instants after stimulation (see colour bar). Color of the lines mark initial PSD size. (B) Absolute change in spine volume after consolidation for different initial PSD areas. (C) Absolute change in PSD areas after consolidation for different initial PSD areas.
